# When do we become more prone to distraction? Progressive evolution of the different components of distractibility from early to late adulthood

**DOI:** 10.1101/2022.08.22.504838

**Authors:** R.S. Hoyer, O. Abdoun, M. Riedinger, R. Bouet, H. Elshafei, A. Bidet-Caulet

## Abstract

Life expectancy has steadily increased for over a century; we thus live longer and are more likely to experience cognitive difficulties such as increased distractibility which can hamper autonomy. This cross-sectional behavioral study aimed to characterize the decline of the cognitive components of distractibility during typical aging, and the onset of this decline. 191 participants from 21 to 86 years old, distributed within seven age groups, were tested using the Competitive Attention Test. Results indicate that cognitive components contributing to distractibility follow different trajectories with aging: voluntary orienting remains stable from 21 to 86 years old, sustained attention decreases while distraction increases between 26 and 86 years old, finally, impulsivity is lower in older compared to younger adults. Increased distractibility in older adults thus seems to result from a dominance of involuntary over voluntary attention processes, whose detrimental effect on performance is partly compensated by enhanced recruitment of motor control.

## INTRODUCTION

Aging is associated with a failure of attention to regulate the processing of irrelevant information (see Healey et al., 2008 for a review). Sub-clinical attention difficulties hamper autonomy and create dependency on others (Ruan et al., 2015). To develop rehabilitation procedures to counteract this loss of autonomy, a prerequisite is to precisely characterize distractibility from early to late adulthood.

“Distractibility” can be conceptualized as a state determining the propensity to have one’s attention captured by irrelevant information, while “distraction” designates the deleterious effect of involuntary attention capture on ongoing behavioral performance. Distractibility relies on a balance between voluntary and involuntary attention mechanisms, allowing to focus while staying alert to surroundings. Voluntary attention promotes the processing of task-relevant stimuli and is internally driven. Involuntary attention is directed by external stimuli and refers to the capture of attention by task-irrelevant, unexpected and salient events, leading to distraction. Beyond attention, behavioral distractibility is shaped by phasic arousal (i.e., phasic alertness) and motor control (whose failure induces impulsivity). Attention, arousal and motor control are underpinned by interconnected brain networks (Aston-Jones & Cohen, 2005; Petersen & Posner, 2012; Posner et al., 1982; Seidler et al., 2010). Distractibility components have been mostly studied separately, through the comparison of two age groups (i.e., younger vs. older adults; Andrés et al., 2006; Brodeur & Enns, 1997; Coyne et al., 1978; Davies & Davies, 1975; ElShafei et al., 2020; Greenwood et al., 1993; Horváth et al., 2009; Iarocci et al., 2009; Jackson & Balota, 2012; Leiva et al., 2014, 2016; Mager et al., 2005; Olk & Kingstone, 2015; Parasuraman et al., 1989; Parmentier & Andrés, 2010). These studies were successful in identifying *what* changes in attention abilities occur between early and late adulthood, but not *when* these changes take place during aging. Furthermore, they yielded inconsistent results: methodologically, this might be explained by the use of broad age ranges within the to-be-compared groups of participants. These contradictory findings are detailed in the followings.

The capacity to orient voluntary attention has been found either unchanged (Greenwood et al., 1993; Iarocci et al., 2009; Olk & Kingstone, 2015) or decreased (Brodeur & Enns, 1997; see also Erel & Levy, 2016 for a review) with aging. The ability to sustain voluntary attention over time (also known as “vigilance”) has been found deteriorating from middle to late adulthood (Berardi, 2001; Davies & Davies, 1975; Fortenbaugh et al., 2015; Jackson & Balota, 2012; Parasuraman et al., 1989; Petton et al., 2019), or to improve with aging (see Vallesi et al., 2021 for a review).

Distraction (quantified as increased reaction times or reduced accuracy to targets preceded by a salient sound; Andrés et al., 2006; Bidet-Caulet et al., 2015; Escera et al., 2000; Näätänen, 1992; Wetzel & Schröger, 2014) was found increased (Andrés et al., 2006; Berti et al., 2013; ElShafei et al., 2020; Leiva et al., 2014, 2016; Parmentier & Andrés, 2010) or unchanged (Horváth et al., 2009; Mager et al., 2005) in older compared to younger adults.

Distractors also trigger a phasic increase in arousal (i.e., alertness) resulting in behavioral benefits in some tasks (Andrés et al., 2006; Bidet-Caulet et al., 2015; Masson & Bidet-Caulet, 2019; Max et al., 2015; Näätänen, 1992; Wetzel et al., 2012). This increase in arousal raises cortical responsiveness via the Locus Coeruleus (Aston-Jones & Cohen, 2005), whose activity is altered with aging (Dahl et al., 2022). However, at the behavioral level, the phasic arousal increase triggered by distractors seems to remain unchanged with aging (Andrés et al., 2006; ElShafei et al., 2020; Parmentier & Andrés, 2010).

Changes in voluntary and involuntary attention, as well as phasic arousal, may urge one to impulsively respond to irrelevant events. Impulsivity is the tendency to act before having fully analyzed a situation, without regard for the consequences of the act to oneself or to others (Barratt & Patton, 1983). Motor inhibition also plays a role in the emergence of impulsive behaviors as it supports the ability to stop an ongoing response. Impulsivity has been found increased (Coyne et al., 1978; Maylor et al., 2011; Nielson et al., 2002) or unchanged (Hong et al., 2014; Hsieh et al., 2016; Lin & Cheng, 2020; Paitel & Nielson, 2021) in older adults compared to younger ones (see also Rey-Mermet & Gade, 2018 for a review).

Using the Competitive Attention Task (CAT; Bidet-Caulet et al., 2015), which provides simultaneous and dissociated measures of several attention facets, recent studies have investigated the brain origins of distractibility in 60-75-year-old adults (ElShafei et al., 2020, 2022). Their magnetoencephalographic findings suggest that elderly have diminished voluntary attention (ElShafei et al., 2020, 2022). In particular, increased distractibility in older adults is associated with reduced top-down inhibition of irrelevant information, a brain mechanism supported by the lateral prefrontal cortex (Amer et al., 2016; Colcombe et al., 2005; (ElShafei et al., 2020b, (2022). From a theoretical perspective, this is aligned with the inhibitory deficit (Hasher & Zacks, 1988) and the frontal aging (West, 1996) hypotheses. However, it remains to elucidate when precisely during aging this prefrontal-related decline starts impacting attention performance.

Using the CAT, the present study aims to outline the evolution of the cognitive components contributing to distractibility from 21 to 86 years, in a large sample of participants (N=191). This paradigm provides behavioral measures of voluntary orienting, sustained attention, distraction, phasic arousal, as well as motor control and impulsivity (Bidet-Caulet et al., 2015; Hoyer et al., 2021). Based on the literature presented above, and in particular on the findings from Elshafei and colleagues (2020, 2022), we hypothesized that, with aging, voluntary orienting, phasic arousal, impulsivity and motor control would remain stable, while sustained attention would progressively decrease and distraction increase, resulting in greater distractibility.

## METHOD

191 subjects from diverse socioeconomic statuses (small employers and own account works, never worked or long term unemployed, managerial and professional occupation, lower supervisory and technical occupations, semi-routine and routine occupations, intermediate occupations, retired; see Fig. S1, Supplemental Material) who spoke fluently French participated in the study. Participants were recruited using e-mail lists and via several senior clubs. They had to fulfill the following inclusion criteria: corrected-to-normal hearing (2 participants wore hearing aid and confirmed that it caused them no discomfort for hearing the different sounds during the experiment), normal or corrected-to-normal vision, no neurological or psychiatric disorders, and no medication affecting the central nervous system taken during the 24 hours preceding the testing session. The samples selected for the study was of convenience: participants were recruited until each of the seven age groups included a minimum of 20 participants (see Tab. 1). Participants were selected to match, as best as possible, the age groups in gender, handedness and education.

**Table 1.** Characteristics of the population sample. Detailed samples, mean age in years, gender, handedness, mean education level (0 = no diploma, 1 = vocational certificate obtained after the 9^th^ grade, 2 = high school diploma; 3 = 12^th^ grade/associate’s degree; 4 = bachelor degree; 5 = master degree and further) and thresholds of auditory perception in dBA by age range (± standard error of the mean, SEM).

| Age range | Number of included participants | Mean age in years | Gender |  | Handedness |  |  | Mean education | Auditory threshold |  |
| --- | --- | --- | --- | --- | --- | --- | --- | --- | --- | --- |
|  |  |  | Female | Male | Right | Left | Ambidextrous |  | Right ear | Left ear |
| 21-25 | 32 | 23.3 ± 0.3 | 50.0 % | 50.0 % | 84.4 % | 15.6 % | 0.0 % | 4.1 ± 0.2 | 25.8 ± 2.2 | 26.7 ± 2.2 |
| 26-30 | 24 | 27.0 ± 0.3 | 58.3 % | 41.7 % | 87.5 % | 8.3 % | 4.2 % | 3.6 ± 0.3 | 31.7 ± 2.5 | 30.9 ± 2.5 |
| 31-40 | 25 | 35.2 ± 0.5 | 60.0 % | 40.0 % | 80.0 % | 20.0 % | 0.0 % | 4.2 ± 0.2 | 31.1 ± 2.1 | 30.6 ± 2.0 |
| 41-50 | 27 | 46.1 ± 0.6 | 74.1 % | 25.9 % | 88.9 % | 7.4 % | 3.7 % | 3.7 ± 0.2 | 28.4 ± 2.4 | 28.7 ± 2.0 |
| 51-60 | 25 | 55.8 ± 0.6 | 56.0 % | 44.0 % | 92.0 % | 4.0 % | 4.0 % | 3.1 ± 0.3 | 32.6 ± 2.5 | 33.6 ± 2.7 |
| 61-70 | 25 | 65.7 ± 0.6 | 60.0 % | 40.0 % | 88.0 % | 0.0 % | 12.0 % | 3.1 ± 0.3 | 35.6 ± 2.5 | 36.2 ± 2.5 |
| 71-86 | 28 | 78.4 ± 0.9 | 85.7 % | 14.3 % | 82.1 % | 7.1 % | 10.7 % | 2.0 ± 0.3 | 42.4 ± 1.4 | 42.8 ± 1.9 |

Data from 5 participants were excluded from the analysis, due to either below-chance performance (correct trial percentage < 50 % in no distractor condition, see Fig. 1: n = 2) or technical issues (n = 3). A total of 186 subjects (86 % right-handed, 9%left-handed, 5 % ambidextrous; 63 % female; 21 to 86 years old) were included in the analysis (see Tab. 1 for details by age ranges). All participants gave written informed consent. This study was conducted according to the Helsinki Declaration, Convention of the Council of Europe on Human Rights and Biomedicine, and the experimental paradigm was approved by the French ethics committee Comité de Protection des Personnes. Note that, to improve readability, “yo” instead of “years-old” or “year-olds” will be used in the Method and Results sections when referring to the participants’ age ranges.

**Figure 1.**
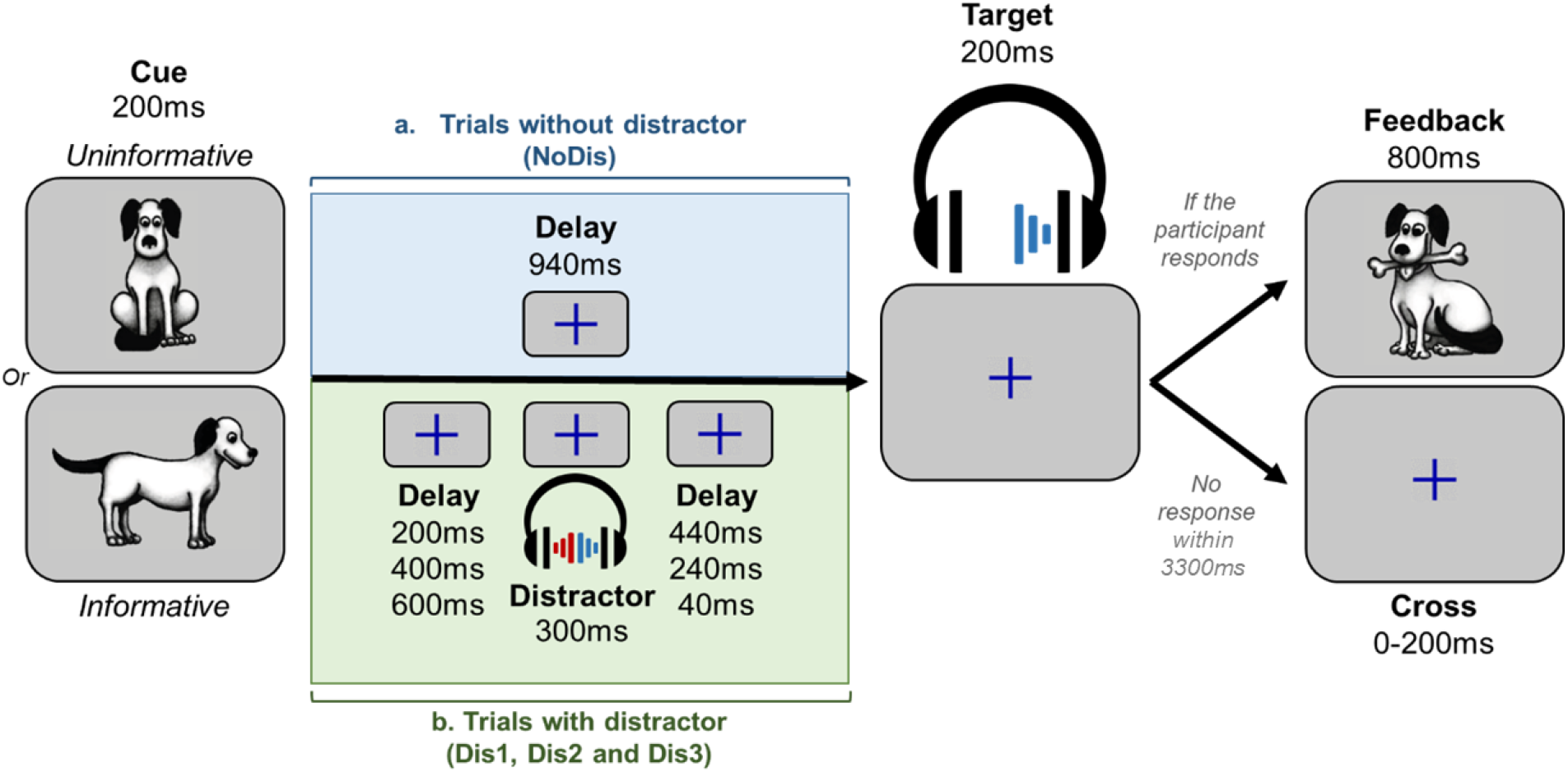
Protocol. *Note*. a) In uninformative trials, a facing-front dog was used as a visual cue, indicating that the target sound would be played in either the left or right ear. In informative trials, a dog visual cue facing left or right indicated in which ear (left or right, respectively) the target sound will be played. If the participant gave a correct answer within the 3300 ms post target offset, feedback (800 ms duration) was displayed. b) In trials with distractors, the task was similar, but a binaural distracting sound - such as a phone ring - was played during the cue-target delay. The distracting sound could equiprobably onsets at three different times: 200 ms, 400 ms, or 600 ms after the cue offset.

### Stimuli and task

A detailed description of the task can be found in a previous study (Hoyer et al., 2021 for more details). In no distractor condition (NoDis; 50 % of the trials), a visual cue (200 ms duration) was followed, after a 940 ms delay, by a target sound (200 ms duration; Fig. 1a). The cue was a dog facing left or right (informative: 75 %), or to the front (uninformative: 25 %). The target sound was a dog bark monaurally presented in headphones. The dog facing left or right (informative) was followed by the taget sound in the left (37.5 %) of right (37.5 %) ear, respectively; the facing-front dog (uninformative) was followed by the target sound in the left (12.5 %) or right (12.5 %) ear. In distractor condition (50 % of the trials), a binaural distracting sound (300 ms duration, 18 different ringing sounds distributed across blocks) was played during the 940 ms delay (Fig. 1b): this sound could be played at three different times - distributed equiprobably - during the delay: 200 ms (Dis1), 400 ms (Dis2) and 600 ms (Dis3) following the cue offset. The target sound was presented at 15 dB SL (around 37.5 dBA) and the distracting sound at 35 dB SL (around 67.5 dBA) in headphones. Cue categories (informative and uninformative) and target categories (left and right) were equally distributed through trials with and without distracting sounds.

Participants were instructed to focus their attention on the cued side, and to press a key as fast as possible when they heard the target sound. Visual feedback (800 ms duration) was displayed when participants detected the target within 3300 ms of its onset, followed by a rest period (inter-trial interval: 1700 ms to 1900 ms). If the participant did not respond in time, the fixation cross was displayed for an additional randomized delay (100 ms to 300 ms).

### Procedure

Participants were tested individually or in small groups (of two or three) in a quiet room. During the task, they were seated in front of a laptop (approximately 50 cm from the screen) that presented pictures and sounds and recorded behavioral responses using Presentation software (Neurobehavioral Systems, Albany, CA, USA). Auditory stimuli were played in headphones. First, the auditory threshold was determined for the target sound, in each ear, for each participant using the Bekesy tracking method. This resulted in an average target threshold across subjects of 32.5 dBA (see Tab. 1 for details by age range). Then, participants received verbal instructions and performed three 4-minutes blocks of 48 pseudo-randomized trials each. The experimental session lasted around 30 minutes.

#### Measure parameters

We used a custom MATLAB program to preprocess behavioral data. The shortest RT for a correct response (RT lower limit) was calculated for every age range (150 ms in the 21 to 60-year-olds and to 200 ms in the 61 to 86-year-olds, see Fig. S2.1, Supplemental Material). For each participant, the longest RT for a correct response (RT upper limit) was calculated from all RT > 0 ms using the Tukey method of leveraging the interquartile range. Based on the lower and upper RT limits, responses can be divided into three categories: (i) responses before the RT lower limit were considered as a part of the false alarm response type; (ii) responses between the lower and the upper RT limits were considered as correct responses; (iii) responses after the RT upper limits were considered as late responses. A total of eight behavioral measures were extracted for each participant (see Tab. 2 and Fig. S2.2, Supplemental Material).

**Table 2.**
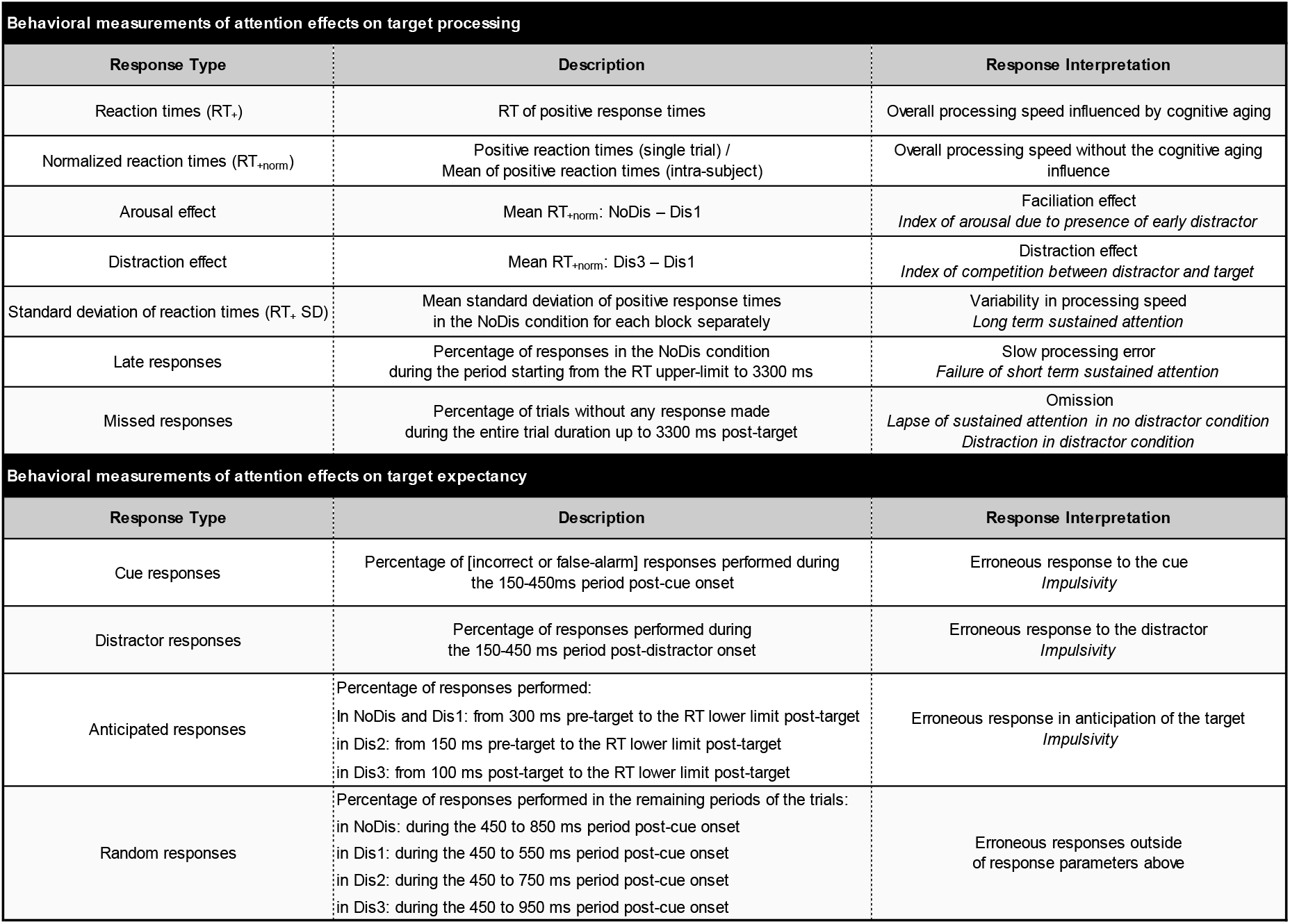
Measure names, detailed criteria for responses categorization in the CAT trials and associated measured constructs.

#### Statistical analysis

To estimate physical tendencies linked to the behavioral measures at the single trial level, we used Generalized Linear Mixed Models (GLMM) in a frequentist approach; models were adapted to the data distribution (i.e., Gaussian for RT related measures, binomial for response types). In order to test the similarity between groups, Bayesian statistics were used. Contrastingly to Frequentist statistics, Bayesian analyses allow to assess the credibility of both the alternative and null hypotheses.

##### Sample characteristics

To confirm that our sample population was similarly distributed across age ranges in block order, gender, handedness and socio-economic statuses (SES), we performed Bayesian contingency table tests. We performed a Bayesian ANOVA on the education level with AGE as between-subject factor to investigate potential differences in education across age ranges. Bayesian statistics were performed using JASP^®^ software (JASP Team, 2021, Version 0.14.1). We reported Bayes Factor (BF_10_) as a measure of evidence in favor of the null hypothesis (BF_10_ 0.33 to 1, 0.1 to 0.33, 0.01 to 0.1 and lower than 0.01: weak, positive, strong and decisive evidence respectively) and in favor of the alternative hypothesis (value of 1 to 3, 3 to 10, 10 to 100 and more than 100: weak, positive, strong and decisive evidence respectively; Lee & Wagenmakers, 2013).

##### Behavioral data analysis

To analyze behavioral data at the single trial level, we used GLMM (Bates et al., 2015). The variability between subjects in raw performance was modeled by defining by-subject random intercepts.

To assess the impact of the manipulated task parameters (cue information and distractor type) and participant age range, on each type of behavioral measure (RT_+_, RT_+_ SD, late responses, missed responses, cue responses, distractor responses, anticipated responses, random responses; see Tab. 2 for more detail), we analyzed the influence of three possible fixed effects and their interaction (unless specified in Tab. 3): the between-subject factor AGE (7 levels: 21-25, 26-30, 31-40, 41-50, 51-60, 61-70 and 71-86); the within-subject factor CUE (2 levels: informative and uninformative); the within-subject factor DISTRACTOR (4 levels: NoDis, Dis1, Dis2 and Dis3). A summary of the data and factors used in statistical modeling can be found in Tab. 3. Because of the different timing for categorizing erroneous responses (see Fig. S2, Supplemental Material), and the low proportion of the cue, random, distractor and late responses, we did not consider the CUE and DISTRACTOR factors in analyses of these measures and only focused on the AGE effect. For anticipated and missed responses, we considered the within-subject factor DISTRACTOR in the analysis as these responses have previously been identified as good markers of distraction and impulsivity in children and young adults (Hoyer et., al. 2021).

**Table 3.**
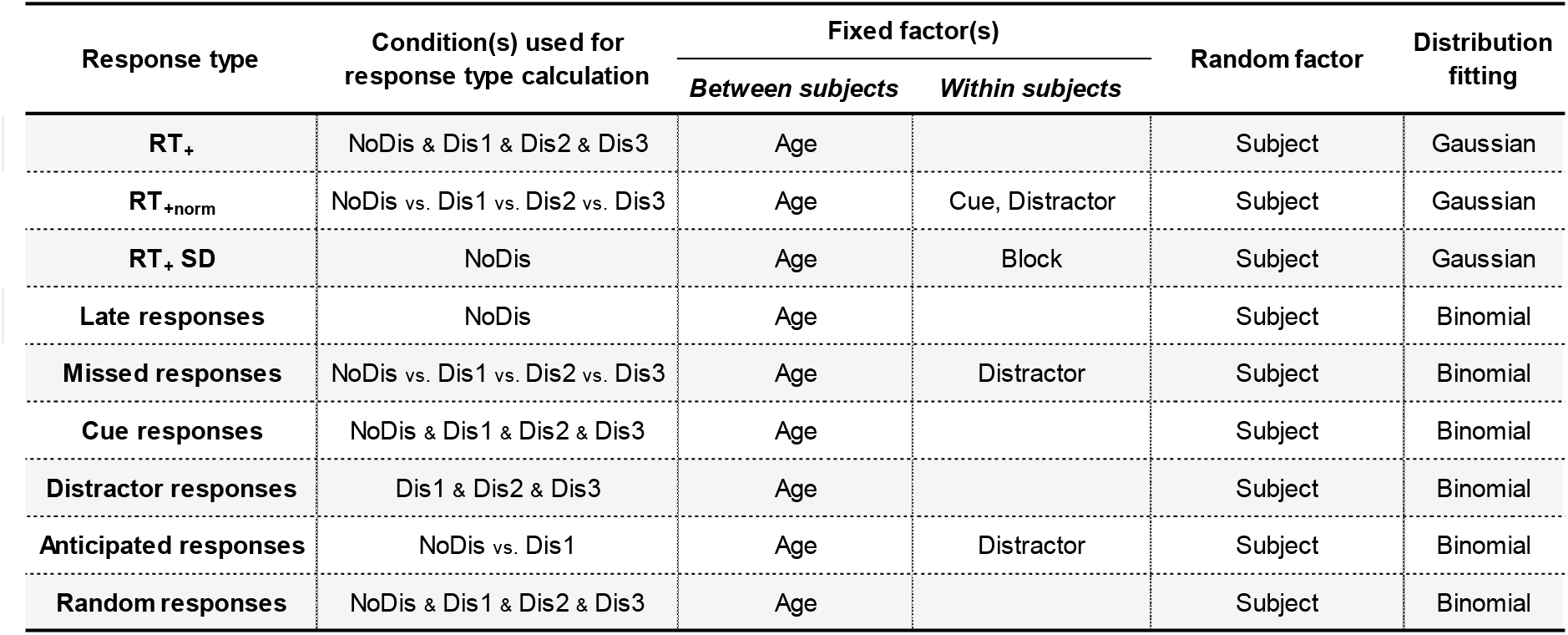
Main statistical analyses according to behavioral response types. *Note*. Experimental conditions, factors and models used as a function of the behavioral measures. Detailed factor levels: DISTRACTOR = NoDis, Dis1, Dis2 and Dis3; CUE = informative, uninformative; Block = first, second and third.

Frequentist statistics were performed in R^®^ 3.4.1 using the lme4 (Bates et al., 2015) and car (Fox & Weisberg, 2018) packages. Both fixed and random factors were considered in statistical modelling. Wald chi-square tests were used for fixed effects in linear mixed-effects models (Fox & Weisberg, 2018). The fixed effect represents the mean effect across all subjects after accounting for variability. We considered results of main analyses significant at p < .05.

When we found a significant main effect or interaction, Post-hoc Honest Significant Difference (HSD) tests were systematically performed using the emmeans package (emmeans version 1.6.3). P-values were considered as significant at *p* < .05 and were adjusted for the number of comparisons performed. In the Results section, we reported the 95 % confidence intervals as a measure of uncertainty; error bars within plots represent quantiles between 5 and 95 %.

###### RT_+_ and RT_+*norm*_

The total average and associated standard error of the mean number of trials with positive RT (RT_+_ et RT_+*norm*_) was of 69.7 ± 0.4 (min: 38, max: 72) in NoDis, 22.3 ± 0.2 (min: 10, max: 24) in Dis1, 22.5 ± 0.2 (min: 12, max: 24) in Dis2 and 22.8 ± 0.2 (min: 12, max: 24) in Dis3 conditions across the overall sample.

To investigate the effect of cognitive aging on the global response speed, raw RT_+_ were fitted to a linear model with AGE only as between subject factor. To avoid analysis bias due to the typical slowing affecting RT during aging (see Leiva et al., 2021 for more details), further analysis were performed on raw RT_+_ normalized at the single trial level using individual mean RT_+_ (at the participant level: RT_+_ single trial / mean RT_+_). Then RT_+*norm*_ were fitted to a linear model, with AGE as between subject factor, and CUE and DISTRACTOR as within subject factors. For post-hoc analysis of the DISTRACTOR by AGE interaction on RT_+*norm*_, we planned to analyze two specific measures of the distractor effect: the distractor occurrence (i.e., the arousal effect; RT_+*norm*_ in NoDis minus RT_+*norm*_ in Dis1) and the distractor position (i.e., the distraction effects; RT_+*norm*_ in Dis3 minus RT_+*norm*_ in Dis1). Based on previous results (Bidet-Caulet et al., 2015; Hoyer et al., 2021; Masson & Bidet-Caulet, 2019), these differences can be respectively considered as good approximations of the facilitation and detrimental distraction effects triggered by distracting sounds (see Fig. 3b).

**Figure 2.**
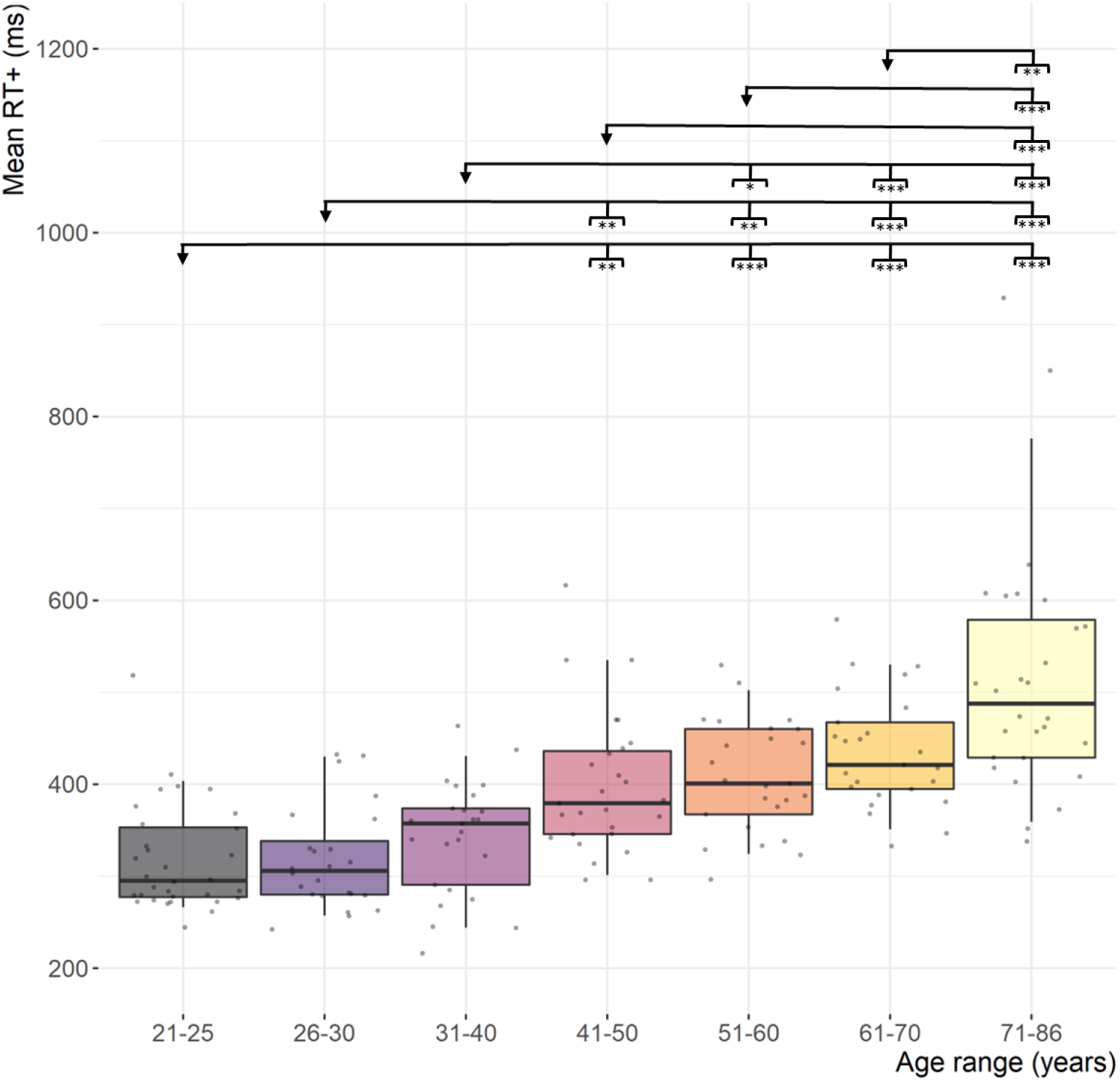
Reaction time according to age. *Note*. Mean reaction time as a function of age range. (*p* < .05 *, *p* < .01 **, *p* < .001 ***). Within each boxplot (Tukey method), the horizontal line represents the group mean, the box delineates the area between the first and third quartiles (interquartile range), the vertical line represents the interval between quantile 5 and 95 (i.e., the dispersion of 90 % of the population); superimposed to each boxplot, the dots represent individual values.

**Figure 3.**
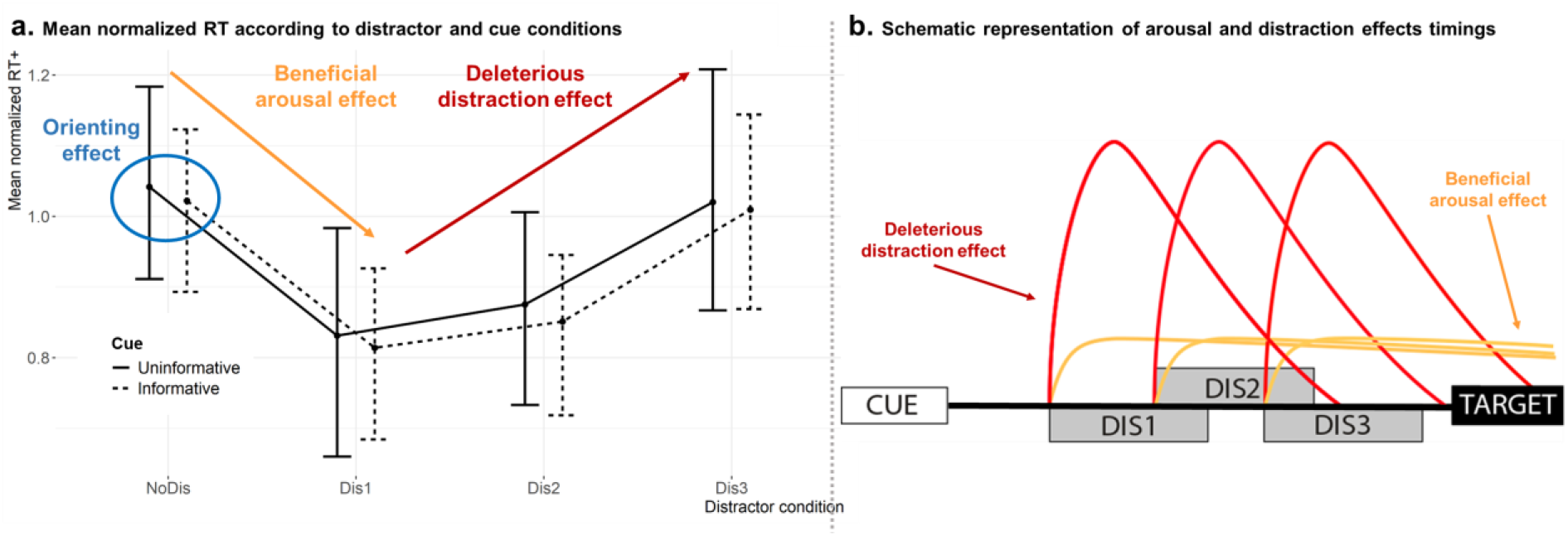
Reaction time according to distractor and cue conditions and schematic representation of the different effects trigger by the distractor according to its timing. *Note. a*) Mean normalized reaction time as a function of the cue (informative or uninformative) and distractor (NoDis, Dis1, Dis2, Dis3) conditions. The error bars represent the interval between quantile 5 and 95 (i.e., the dispersion of 90 % of the population). b) Schematic representation of the facilitation arousal effect (yellow lines) and the distraction effect (red lines) timing during the CAT trials.

###### Other measures

RT_+_ SD were log-transformed at the single trial scale to be better fitted to a linear model with Gaussian family distribution, with the fixed factors AGE and GENDER as between-subject factor and BLOCK (3 levels) as within subject factor.

Response types were fitted to a linear model with binomial distribution without transformation. Missed responses were fitted to a linear model with AGE as between-subject factor and DISTRACTOR as within subject factor (see Tab. 3). Late, cue, random and distractor responses were fitted to a linear model with AGE as between-subject factor (see Tab. 3). Anticipated responses were fitted to a linear model with fixed factors AGE as between-subject factor and DISTRACTOR as within subject factor (see Tab. 3). Because of the important differences in the duration of the anticipated responses windows between distractor conditions (see Fig. S2, Supplemental Material), the GLMM was performed on the NoDis and Dis1 conditions, only (same timeframe for anticipated responses in these two conditions).

## RESULTS

### Population characteristics

Using Bayesian contingency table tests, we found a positive evidence for a similar distribution in gender (BF_10_ = 0.103) across age ranges. We also observed a decisive evidence for a similar distribution in handedness (BF_10_ = 2.057^e^-6) and block order (BF_10_ = 1.086^e^-8) across age ranges. By contrast, we found decisive evidence for a non-uniform distribution in SES characteristics (BF_10_ = 5.707^e^+25), the youngest participants (21-25yo) being mostly students and the oldest ones (71-86yo) being mostly retired (see Fig. S1, Supplemental Material). The Bayesian ANOVA carried out on education level showed that mean education decreases with age (for detailed post-hoc contrasts see Tab. S3, Supplemental Material). This cutback in education level with age can be explained by the fact that access to graduate studies has been made easier these last decades (e.g., construction of universities, availability of fellowships).

### RT_+_

As expected, we observed a main effect of the AGE on RT_+_ (*χ*^2^ (6) = 141.0; *p* < .001; Fig. 2a). Tukey post-hoc analysis indicated that the 21 to 30yo were faster than the 41 to 86yo, the 31-40yo were faster than the 51-86yo and, finally, the 51 to 70yo were faster than the 71-86yo (see Fig. 2 and Tab. 4 for differences, confidence intervals and *p* values). RT_+_ thus progressively increases from 31 to 86 years of age.

**Table 4.** Details for post-hoc analyses of RT_+_.

| Age ranges<br>*A > or < B* | Estimate | Confidence<br>intervals<br>*[Lower, Upper]* | *P* value |
| --- | --- | --- | --- |
| 21-25 < 41-50 | -80.1 ms | [-140.2, -20.0] | $p = .002$ |
| 21-25 < 51-60 | -89.8 ms | [-151.2, -28.4] | $p < .001$ |
| 21-25 < 61-70 | -117.4 ms | [178.8, -56.0] | $p < .001$ |
| 21-25 < 71-86 | -198.2 ms | [-257.8, -138.7] | $p < .001$ |
| 26-30 < 41-50 | -80.3 ms | [-144.9, -15.8] | $p = .005$ |
| 26-30 < 51-60 | -90.0 ms | [-155.7, -24.2] | $p = .001$ |
| 26-30 < 61-70 | -117.6 ms | [-183.3, -51.9] | $p < .001$ |
| 26-30 < 71-86 | -198.2 ms | [-134.4, -262.4] | $p < .001$ |
| 31-40 < 51-60 | -65.9 ms | [-131.0, -0.8] | $p = .045$ |
| 31-40 < 61-70 | -93.5 ms | [-158.6, -28.5] | $p < .001$ |
| 31-40 < 71-86 | -174.4 ms | [-237.6, -111.0] | $p < .001$ |
| 41-50 < 71-86 | -118.1 ms | [-180.2, -56.1] | $p < .001$ |
| 51-60 < 71-86 | -108.5 ms | [-171.8, -45.1] | $p < .001$ |
| 61-70 < 71-86 | -80.8 ms | [-144.1, -17.5] | $p = .003$ |

### RT_+*norm*_

A main effect of CUE (*χ*^2^ (1) = 10.34; *p* = .002; Fig. 3a) on RT_+*norm*_ was observed, indicating that, irrespective of age, participants were faster when the cue was informative (1.000, CI = [0.664, 1.010]) rather than uninformative (1.010, CI = [0.994, 1.030]). The interaction CUE by AGE was found non-significant (*χ*^2^ (6) = 8.79; *p* = .186; Fig. 3a).

As expected, we did not observe a main effect of AGE on RT_+*norm*_ (*χ*^2^ (6) = 0.06;*p* = 1.000), suggesting that the normalization method we used was appropriate. The main effect of DISTRACTOR on RT_+*norm*_ was significant (*χ*^2^ (3) = 2651.87.;*p* < .001; Fig. 3a). An AGE by DISTRACTOR interaction was also significant (*χ*^2^ (18) = 69.52; *p* < .001). Planned post-hoc contrasts were carried on the different RT effects triggered by the distractor occurrence (see Method section and Fig 4b and 4c for more details). Post-hoc indicated that the 21-25yo presents an increased NoDis-Dis1 arousal effect compared to the 26-30yo, 41-50yo and the 51-60yo; the 71-86yo also showed an increased arousal effect compared to the 41-50yo and the 51-60yo (see Fig. 4b and Tab. 5 for differences, confidence intervals and *p* values). In addition, the Dis3-Dis1 distraction effect was increased in the 21-25yo compared to the 26-30yo, in the 41-86yo compared to the 26-30yo and in the 61-86yo compared to the 31-40yo (see Fig. 4c and Tab. 5 for differences, confidence intervals and *p* values). Overall, RT_+*norm*_ analysis indicates that the arousal facilitation effect decreases after 25yo and increases after 60yo, while the distraction effect progressively increases from 26 to 86 years of age.

**Figure 4.**
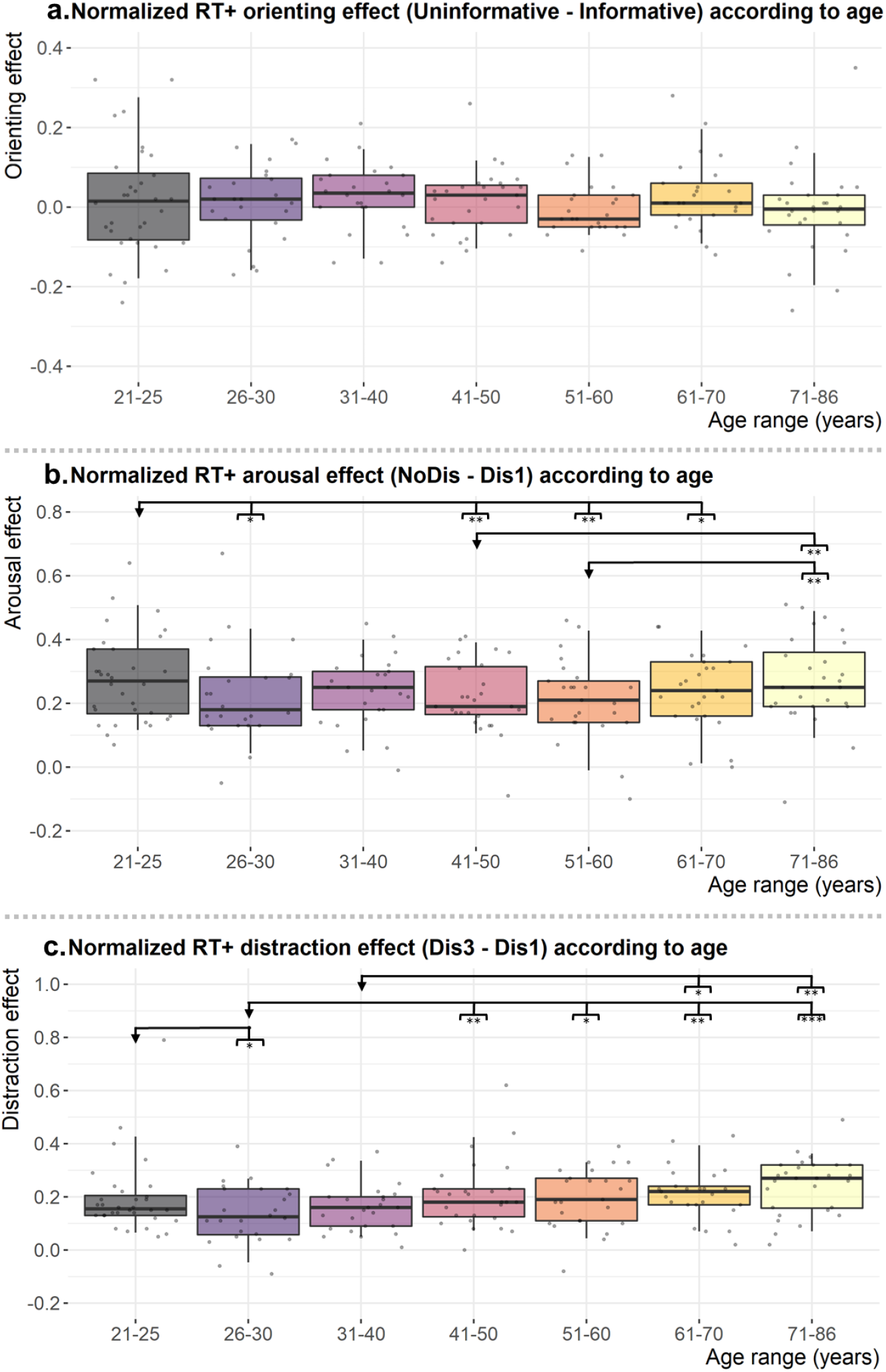
Normalized reaction time orienting, arousal and distraction effects according to age. *Note*. a. Normalized reaction time NoDis Uninformative – NoDis Informative orienting effect as a function of age range. b. Normalized reaction time NoDis-Dis1 arousal effect as a function of age range. c. Normalized reaction time Dis3-Dis1 distraction effect as a function of age range. (*p* < .05 *, *p* < .01 **, *p* < .001 ***). Before calculating differences, RT_+_ have been normalized as follow: participant’s RT_+_ single trial / participant’s mean RT_+_. Within each boxplot (Tukey method), the horizontal line represents the group mean, the box delineates the area between the first and third quartiles (interquartile range); the vertical line represents the interval between quantile 5 and 95 (i.e., the dispersion of 90 % of the population); superimposed to each boxplot, the dots represent individual values.

**Table 5.** Details for significant results of planned post-hoc analyses of normalized RT_+_.

|  | Age ranges<br>*A > or < B* | Estimate | Confidence<br>intervals<br>*[Lower, Upper]* | *P* value |
| --- | --- | --- | --- | --- |
| **Arousal effect**<br>**NoDis-Dis1** | 21-25 > 26-30 | 0.049 | [0.008, 0.089] | $p = .018$ |
| | 21-25 > 41-50 | 0.054 | [0.016, 0.010] | $p = .006$ |
| | 21-25 > 51-60 | 0.066 | [0.026, 0.106] | $p < .001$ |
| | 41-50 < 71-86 | -0.047 | [-0.087, 0.006] | $p = .024$ |
| | 51-60 < 71-86 | -0.058 | [-0.100, -0.016] | $p = .006$ |
| **Distraction effect**<br>**Dis3-Dis1** | 21-25 > 26-30 | 0.087 | [-0.008, 0.106] | $p = .023$ |
| | 26-30 < 41 50 | -0.067 | [-0.119, -0.016] | $p = .010$ |
| | 26-30 < 51-60 | -0.058 | [-0.111, -0.006] | $p = .030$ |
| | 26-30 < 61-70 | -0.075 | [-0.127, -0.024] | $p = .004$ |
| | 26-30 < 71-86 | -0.104 | [-0.154, -0.052] | $p < .001$ |
| | 31-40 < 61-70 | -0.054 | [-0.105, -0.002] | $p = .040$ |
| | 31-40 < 71-86 | -0.082 | [-0.133, -0.031] | $p = .002$ |

### RT_+_ SD

RT_+_ SD was modulated by the AGE (*χ*^2^ (6) = 140.00; *p* < .001; Fig. 5): HSD post-hoc comparisons indicated that the 41 to 86yo had an increased RT variability compared to the 21 to 30yo; the 51 to 86yo had more variable RT_+_ than the 31-40yo; finally, RT_+_ SD was larger in the 71-86yo compared to the 41-70yo participants (see Tab. 6 for differences, confidence intervals and *p* values). We also observed a main effect of the BLOCK on RT_+_ SD (*χ*^2^ (2) = 104.01; *p* < .001): according to HSD post-hoc comparisons, participants had less variable RT_+_ in the third compared to the first (difference (diff.) = −42.3 ms, CI = [−50.6, −34.0]; *p* < .001) and second (diff. = −14.6 ms, CI = [−23.0, −6.4];*p* < .001) blocks, as well as during the second block compared to the first one (diff. = −27.6 ms, CI = [−36.0, −19.4]; *p* < .001). There was no AGE by BLOCK interaction (*χ*^2^ (12) = 5.50; *p* = .942).

**Figure 5.**
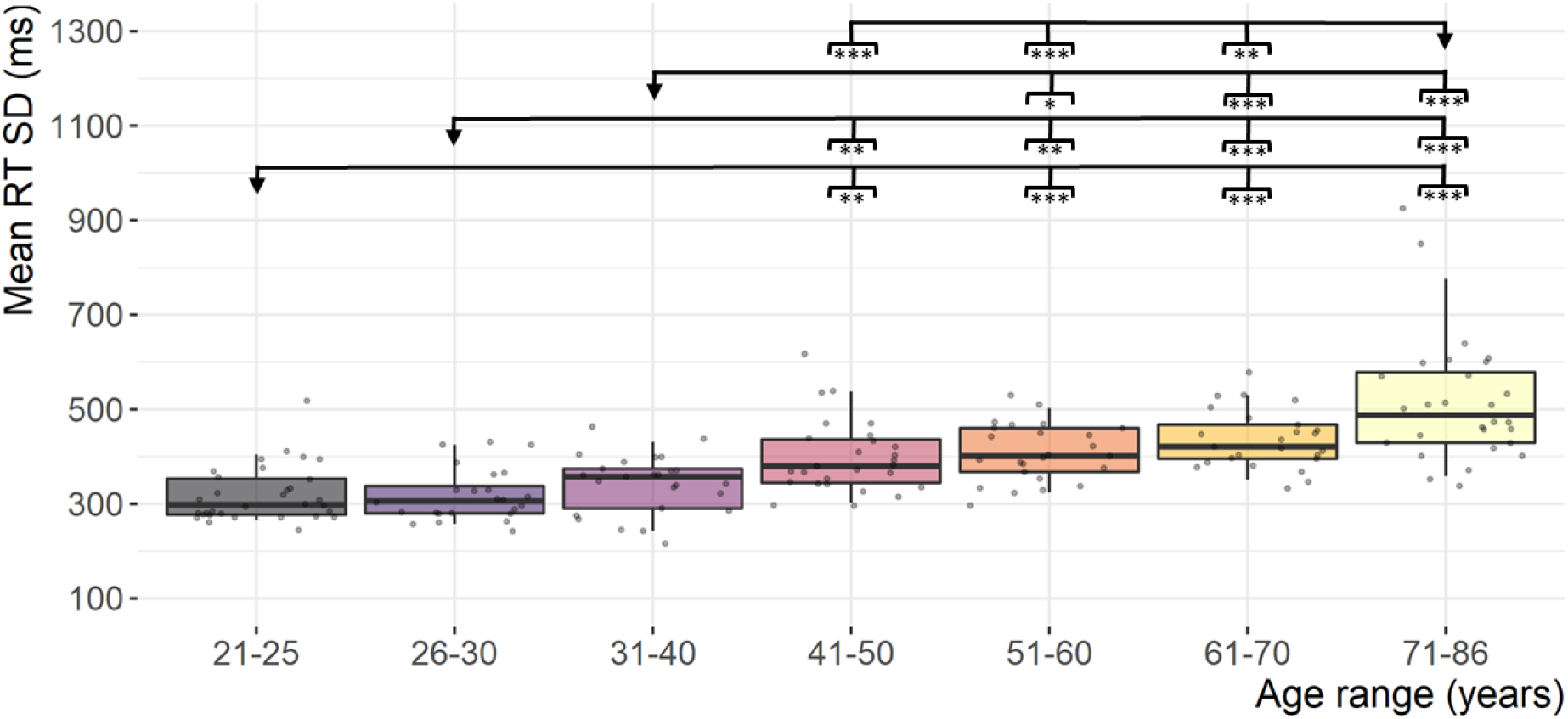
Reaction time variability according to age. *Note*. Reaction time standard deviation (RT_+_ SD) across blocks as a function of age range. (*p* < .05 *, *p* < .01 **, *p* < .001 ***). Within each boxplot (Tukey method), the horizontal line represents the group mean, the box delineates the area between the first and third quartiles (interquartile range the vertical line represents the interval between quantile 5 and 95 (i.e., the dispersion of 90 % of the population); superimposed to each boxplot, the dots represent individual values.

**Table 6.** Details for significant results of post-hoc analyses of RT_+_ SD.

| Age ranges<br>*A > or < B* | Estimate | Confidence<br>intervals<br>*[Lower, Upper]* | *P* value |
| --- | --- | --- | --- |
| 21-25 < 41-50 | -79.5 ms | [-140.2, -18.8] | $p = .003$ |
| 21-25 < 51-60 | -89.4 ms | [-151.4, -27.5] | $p < .001$ |
| 21-25 < 61-70 | -116.6 ms | [-178.5, -54.6] | $p < .001$ |
| 21-25 < 71-86 | -196.8 ms | [-256.9, -136.7] | $p < .001$ |
| 26-30 < 41-50 | -80.7 ms | [-145.9, -15.6] | $p = .005$ |
| 26-30 < 51-60 | -90.7 ms | [-157.0, -24.3] | $p = .001$ |
| 26-30 < 61-70 | -117.8 ms | [-184.2, -51.5] | $p < .001$ |
| 26-30 < 71-86 | -198.0 ms | [-262.6, -133.4] | $p < .001$ |
| 31-40 < 51-60 | -66.2 ms | [-131.9, -0.6] | $p = .047$ |
| 31-40 < 61-70 | -93.37 ms | [-159.0, -27.7] | $p < .001$ |
| 31-40 < 71-86 | -173.6 ms | [-237.5, -109.7] | $p < .001$ |
| 41-50 < 71-86 | -117.3 ms | [-179.9, -54.7] | $p < .001$ |
| 51-60 < 71-86 | -107.4 ms | [-171.3, -43.485] | $p < .001$ |
| 61-70 < 71-86 | -80.2 ms | [-144.1, -16.338] | $p = .005$ |

To sum up, RT_+_ SD progressively increases from 31 to 86 years of age. Furthermore, irrespective of the age, RT_+_ SD gradually reduces across the three blocks of the experiment.

### Global accuracy

The proportion of the different types of behavioral responses according to age is depicted in Fig. 6. The average correct response rate was 87.8 %). No main effect of AGE was found for cue responses (cue responses, total average: 0.2 %), distractor responses (distractor responses, total average: 2.5 %), random responses (random responses, total average: 0.1 %), and late responses (late responses, total average: 10.3 %).

**Figure 6.**
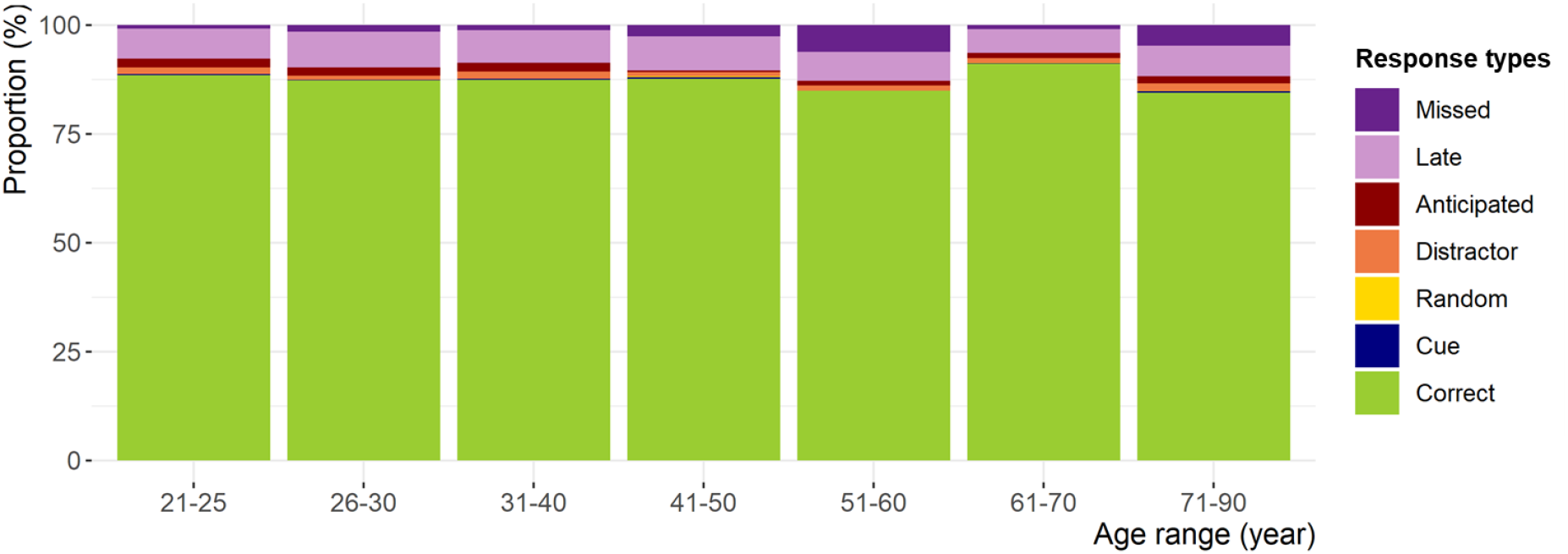
Response type proportions according to age.

### Missed responses

The rate of missed responses (missed responses, total average: 2.4 %) was modulated by AGE (*χ*^2^ (6) = 20.42; *p* = .002; Fig. 7a). HSD post-hoc comparisons revealed that 51-70 and 71-86yo participants missed more the target than the 61-70yo (respectively: diff. = −0.1 units, CI = [-1.7, 1.6], and diff. = −0.7, CI = [-2.1, 0.8], respectively).

**Figure 7.**
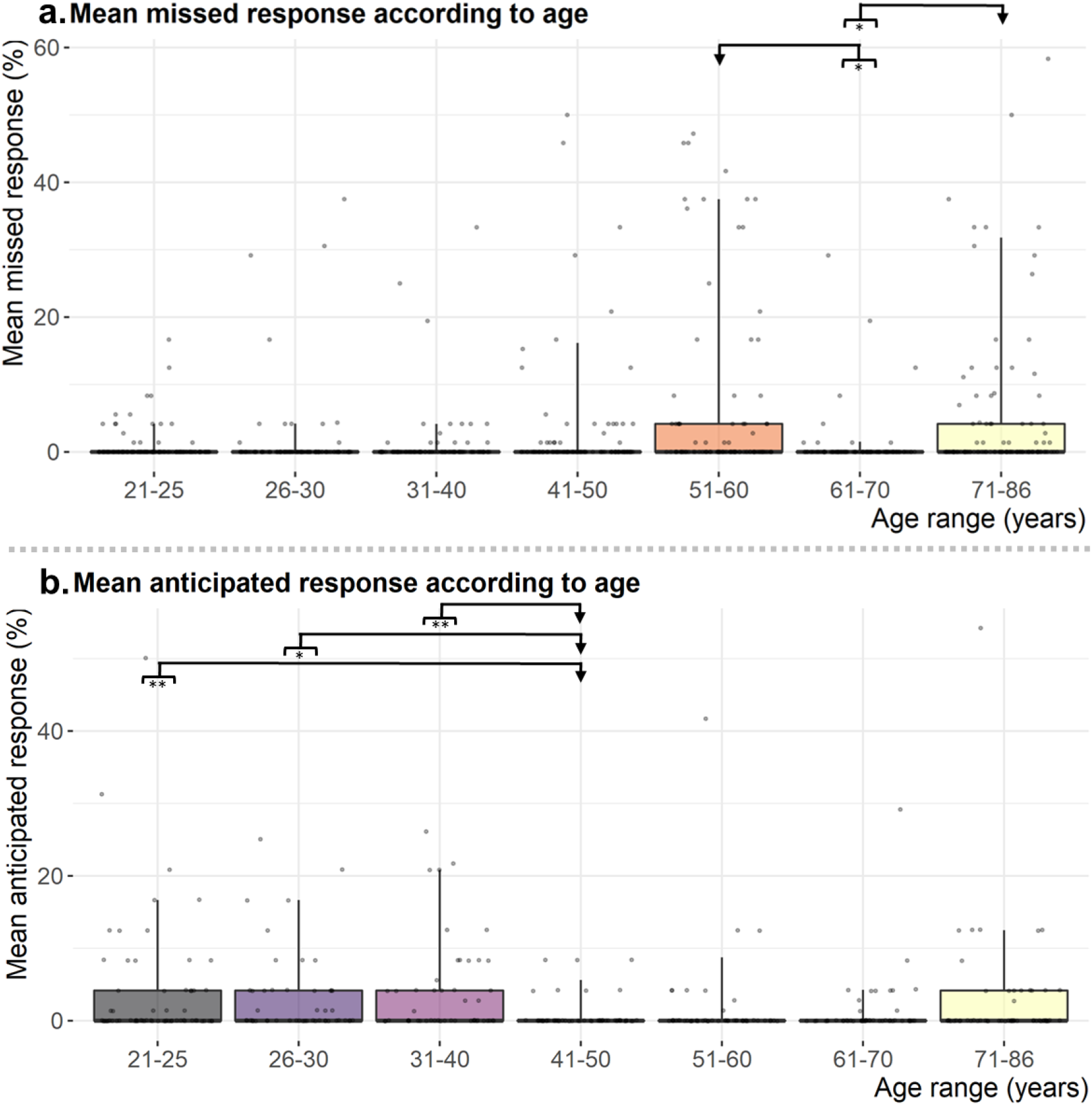
Response type proportions according to age. *Note*. a) Missed responses percentage as a function of the age range. b) Anticipated responses percentage (NoDis and Dis1) as function of the age range. (p < .05 *, p < .01 **). Within each boxplot (Tukey method), the horizontal line represents the group mean, the box delineates the area between the first and third quartiles (interquartile range); the vertical line represents the interval between quantile 5 and 90 (i.e., the dispersion of 95 % of the population); superimposed to each boxplot, the dots represent individual values.

Thus, the missed response rate is greater in participants of 51-60 and 71-86 years of age.

### Anticipated responses

The rate of anticipated responses (anticipated responses, total average: 2.6 %) was modulated by AGE (*χ*^2^ (6) = 24.61; *p* < .001; Fig. 7b). Post-hoc HSD analysis showed that the 41-50yo (0.4 units, i.e., 0.8 %) made less anticipated responses than the 21-25yo (diff. = −1.8, CI = [-2.8, 0.8]), 26-30yo (diff. = −1.7, CI = [-2.8, 0.7]) and 31-40yo (diff. = −2.0, CI = [−3.0, −0.9]). We also observed a main effect of the DISTRACTOR on the anticipated responses rate (*χ*^2^ (1) = 264.80; *p* < .001) indicating that participants anticipated the target much more in Dis1 compared to the NoDis condition (diff. = −3.4, CI = [−3.8, −3.0]).

In summary, the anticipated response rate is larger in participants aged from 21 to 40 years. Furthermore, irrespective of the age, more anticipated responses are observed in trials with early distractors.

## DISCUSSION

The present cross-sectional study provides, for the first time, simultaneous but distinct measures reflecting the evolution of the distractibility components from 21 to 86 years old (7 age ranges). These findings reveal the paths taken by voluntary orienting, sustained attention, distraction, phasic arousal, as well as impulsivity and motor control alongside aging.

Voluntary attention orienting does not significantly change through age, suggesting a preserved ability to orient attention toward relevant targets in elderly (Greenwood et al., 1993; Iarocci et al., 2009; Olk & Kingstone, 2015). In line with studies showing a decline in sustained attention in older adults (Berardi, 2001; Davies & Davies, 1975; Fortenbaugh et al., 2015; Jackson & Balota, 2012; Parasuraman et al., 1989; Petton et al., 2019), we found that RT variability in no distractor trials increases from 26 years old to the older age.

The present findings confirm that distraction is increased in elderly (Andrés et al., 2006; Berti et al., 2013; ElShafei et al., 2020; Leiva et al., 2014, 2016; Parmentier & Andrés, 2010), but also reveals that this heightened distraction developed gradually from early to late adulthood. Increase distraction effect after age 60 is assumed to ensue from greater brain processing of distractors, reduced recruitment of frontal-mediated inhibitory mechanisms, as well as prolonged reorientation towards the task (ElShafei et al., 2020, 2022; Horváth et al., 2009; Mager et al., 2005).

Distractors can also trigger a transient increase in phasic arousal (i.e., in alertness) and speed the response to a subsequent target (Aston-Jones & Cohen, 2005). Here, the arousal-related facilitation effect is greater in participants of 21-25 and 71-86 years of age, suggesting that phasic arousal first decreases during early adulthood and then rebounds in old age. Previous studies have found that the facilitation effect was not modulated by age (Andrés et al., 2006; ElShafei et al., 2020; Parmentier & Andrés, 2010), but they only compared younger and older adults from wide age ranges (18-29 and 50-83 years old). This method is not appropriate to apprehend the subtle attention changes which occur with aging.

In the present study, impulsivity was assessed from different behavioral measures provided by the CAT (cue, random, distractor and anticipated responses). Among these measures, only the proportion of anticipated responses changes with age: adults aged 21-40 are more likely to impulsively press the button before target than their older peers, suggesting that motor inhibition is increased after 40-years-old.

In line with both the frontal aging (Greenwood, 2000; West, 1996) and the inhibitory deficit (Hasher & Zacks, 1988) theories, this study shows that increased distractibility with aging originates from reduced sustained attention and increased distraction. The frontal lobes decline with age has been related to weakened sustained attention abilities (Vallesi et al., 2021). Enhanced distraction can stem from an inhibitory deficit: lessening distractor processing in the brain would become increasingly laborious with age (ElShafei et al., 2020, 2022). Our findings suggest that both reduction in sustained attention and enhancement in distraction start around 40 years old.

Most of the present findings are aligned with the prominent theories on the aging brain, whereby cognitive processes supported by frontal functions decline with age. Voluntary attention orienting seems however preserved with aging while relying on the frontal lobe integrity (e.g., Bidet-Caulet et al., 2015). An explanation is the development of compensatory mechanisms. Stronger engagement of motor regions and reduced inhibition of irrelevant brain areas has been observed during target expectancy in participants aged of 61 to 75 years while performing the CAT (ElShafei et al., 2020, 2022). The compensation of lower attention efficiency by higher motor preparation in elderly might result in preserved voluntary attention orienting performance. One may have expected that larger activation of motor preparation processes before target occurrence would lead to more anticipated responses in elderly, but the opposite trend was observed in this study: after 40 years of age, anticipated responses progressively fade. This reduction in impulsivity may actually be related to the general slowdown in response times which has been consistently observed in older adults, and is likely to ensue from the age-related decline of the central and peripheral motor systems (see Seidler et al., 2010 for a review) and a decrease in tonic arousal (Dahl et al., 2022; Lee et al., 2018; Mather & Harley, 2016; Müller-Oehring et al., 2013). The heightening of phasic arousal after 70 years of age may also be a compensatory mechanism in late adulthood to alleviate the diminishing efficiency of its tonic counterpart (Dahl et al., 2022), but also to compensate behavioral cost related to increased distraction (Gallant et al., 2020).

Compensatory strategies are at the heart of the scaffolding theory of aging and cognition (STAC). This theory assumes that two kinds of brain plasticity occur throughout aging: a negative (leading to cognitive decline) and a positive (supporting compensatory mechanisms) one (Goh & Park, 2009; Reuter-Lorenz & Park, 2014). Compensatory mechanisms can be established during aging to cope with enhanced distractibility. Motor preparation might compensate for reduced voluntary attention abilities such as orienting, but would not be sufficient to compensate the decline of processes requiring more cognitive resources, such as sustained attention. In late adulthood, the enhancement of motor preparation might also reduce impulsivity, and the phasic arousal increase might compensate for enhanced distraction and lower tonic arousal.

### Conclusions

The present behavioral study shows that the different cognitive components contributing to distractibility follow different trajectories from early to late adulthood: voluntary orienting is stable from 21 to 86 years of age, sustained attention progressively decreases, distraction increases between 26 and 86 years old, phasic arousal decreases after 25 years of age but upturns after 70 years old and, eventually, impulsivity is greater in 21- to 40-year-old than in older adults. Taken together, these findings suggest that increased distractibility in older adults ensues from an attentional imbalance characterized by a dominance of involuntary over voluntary attention processes. By shedding light on the nonlinear evolutive characteristics of the distractibility components from early to late adulthood, the present findings further emphasize the relevance of using several restrained age ranges in cross-sectional studies of aging.

## CONTEXT

Distractibility relies on different cognitive facets, whose evolution with aging has, so far, remained poorly understood. In a large sample of participants (N=423), we previously used the CAT to delineate the developmental trajectories of several distractibility components from childhood to early adulthood (Hoyer et al., 2021, 2022). The present study shed light on the simultaneous, but distinct, - trajectories taken by the distractibility components from early to late adulthood. Attention Deficit/Hyperactivity Disorder (ADHD) in older adults (Michielsen et al., 2012) is often misdiagnosed, and can even be mistaken with dementia (Callahan et al., 2022; Sasaki et al., 2022). Thus, the present findings could help to dissociate sub-clinical and pathological attention difficulties in medical settings. With this in mind, we are currently performing studies to show the content and criterion validity of the CAT, to further enable its use in clinical settings.

## AKNOWLEDGMENTS

We thank J. Hemmerlin, M. Navarro and H. Drissi for their help in recruiting and testing participants. We also would like to thanks all participants. The CAT application is protected by a trademark “CoLeT”.

